# Comparative Mutagenesis of SARS-CoV-2 Nonstructural Proteins (NSPs) Across Variants: The Case for RdRp as a Therapeutic Target

**DOI:** 10.1101/2022.10.15.512346

**Authors:** Nathan Lanclos

**Affiliations:** University of South Florida

## Abstract

Severe acute respiratory syndrome coronavirus 2 (SARS-CoV-2) pathogenicity has been studied extensively from the perspective of structural (S, E, M, N) proteins for purposes in vaccine development. The virus’ nonstructural protein (nsp) components are less characterized, and demonstrate significant potential in efforts to develop novel therapeutic agents. NSP 7, 8, and 12, formed from the cleavage of pp1a and pp1ab polyproteins, comprise the viral replicase (RdRp) complex^1^, the site for the mechanism of action of Remdesivir^2^. Presented herein is a phylogenetic analysis for the evolution of SARS-CoV-2 replicase components between variant and related coronaviruses with the aim to delineate its current and long-term efficacy as a drug target.

## Introduction

SARS-CoV-2 is a type of coronavirus, under the family *Coronaviridae*, demonstrating significant similarity to known species in genus *Betacoronavirus*, specifically SARS-CoV and MERS^**3**^, which have both independently been the cause of other significant outbreaks in the last twenty years. These classifications of virus express broad range virulence for hosts, with many historically, SARS-CoV-2 included, being of zoonotic origin. An article in early 2020 depicted the virus as an emerging threat, isolated from a seafood market in the Wuhan province of China^**4**^. Wuhan Hu-1 has a genome length of 29903bp, and demonstrates a characteristic ribosomal frameshift mechanism essential to the expression of polyproteins in RNA based coronaviruses.

SARS-CoV-2’s genome is comprised of two major open reading frames, ORF1a and ORF1ab, which code for nonstructural proteins 1-16 and represent about 75% of the genome (∼20kb). The genome also contains genes for four structural proteins (spike (S), envelope (E), membrane (M), and nucleocapsid (N)), as well as 11 other accessory proteins (3a, 3b, 3c, 3d, 6, 7a, 7b, 8, 9b, 9c, 10) ^**5**^. These accessory proteins are mostly non-essential to viral replication, and have been implicated with numerous cellular interactions, namely interferon antagonism and immune evasion, but are yet to have clearly elucidated functions^**5**^. Structural proteins of the virus, located at the 3’ end of the genome, are well characterized and contribute heavily to viral stability and host interactions. Spike (S) protein, found on the outside of the nucleocapsid, contains the receptor binding domain that allows the virus to bind host ACE2, and has been subject to major scientific efforts in vaccine development. Other structural proteins E, M, and N have other roles in pathogenesis, and are generally well conserved across coronaviruses^**6**^.

Here we turn our attention to nonstructural proteins, of which have varying degrees of importance in regulating cell activity and replication. Functional analysis of these proteins is not significantly explored, with many of the sources for contemporary knowledge on them dating back to research following SARS in 2003. NSP 1 suppresses host immune function^**7**^, NSP 2 and 3 are not well understood, but may be involved with cellular signaling pathways^**8-9**^. NSP 4 is thought to play a role in the secretory pathway for the replication complex^**10**^, NSP 5 is a “3C-Like Protease” that processes cleavage sites during viral replication^**11**^. NSP 6 from avian coronaviruses produces autophagosomes, but is yet to be characterized in the context of SARS-CoV-2^**12**^. NSP 7 and 8 are co-factors to the viral RNA polymerase with undefined functions^**13**^. NSP 9 is known to have some function in viral replication^**13**^, NSP 10 is a cofactor in the methyltransferase complex^**14**^, and NSP 11 function is unknown^**15**^. NSP 12 is the primary RNA polymerase in viral replication^**13**^, NSP 13 is the viral helicase, it might have 5’ cap activity and has demonstrated enhanced function while bound to NSP 12^**16**^. NSP 14 methylates the 5’ RNA cap with NSP 16, whose activities are enhanced by NSP 10^**15**^. NSP 15 is a uridine endoribonuclease for host immune evasion^**17**^. A summary of these nonstructural proteins are shown in Table 1. below.

**Table 1.** Coronavirus NSP characterization

| Protein | Understood Function | Characterized | Reference # |
| --- | --- | --- | --- |
| NSP 1 | host immune evasion | yes | 7 |
| NSP 2 | cell signaling | no | 8,9 |
| NSP 3 | cell signaling | no | 8,9 |
| NSP 4 | replication signaling | no | 10 |
| NSP 5 | protease | yes | 11 |
| NSP 6 | autophagy | no | 12 |
| NSP 7 | replicase complex cofactor | partially | 13 |
| NSP 8 | replicase complex cofactor | partially | 13 |
| NSP 9 | viral replication | no | 14 |
| NSP 10 | methyltransferase cofactor | yes | 15 |
| NSP 11 | no known function | no | 16 |
| NSP 12 | RNA dependent polymerase | partially | 13 |
| NSP 13 | helicase | partially | 17 |
| NSP 14 | 5' methylation | partially | 18 |
| NSP 15 | uridine endoribonuclease | partially | 19 |
| NSP 16 | 5' methylation | partially | 18 |

The gaps in knowledge on the virus pose a vulnerability in the effort to develop therapeutics and vaccines to fight the spread of the virus, evidenced by the increased mutagenesis of variants that began circulating after Alpha Pango lineages dispersed out of southeastern Asia, and the lack of knowledge on mechanisms of selection pushing its evolution. In particular, the RNA dependent RNA polymerase (RdRp) complex of SARS-CoV-2 is of interest as a drug target for its function as the viral replicase. Out of the 16 non-structural proteins described, the primary complex is composed of NSP 7, 8, and 12, with evidence of other interactions with NSP 4, 9, and 13 (Table 1.). As of late 2021, the only FDA approved treatment for SARS-CoV-2 is Remdesivir (although reports are now surfacing that there might be unaccounted side effects^**18**^), an antiviral produced by Gilead, with production starting as early as 2009 as a treatment for RSV^**19**^. Remdesivir’s mode of action from a molecular perspective involves the drug acting as a nucleoside analog that stalls the replicase complex via a translocation barrier during the replication process^**20**^.

Remdesivir, as well as other antivirals being tested, take advantage of the importance of the replicase in viral proliferation, but raises the question as to the effects of adding a positive selection pressure to the mutagenesis of the polymerase. Literature on the evolution of the virus has suggested that the use of polymerase inhibiting therapeutics has the potential to increase mutagenicity of the virus and promote the evolution of unfavorable genes^**21**^. Other studies have evaluated the effects of specific mutations, in particular, P4715L and P323L (both NSP 12 mutants), and established that variants with these mutations have thus far emerged to be the most epidemiologically successful variants^**22**^. To this end, this work aims to elucidate the connection with mutations in the core replicase and associated nonstructural proteins to the evolution of SARS-CoV-2 variants, and infer on how pathogenicity has been affected downstream of these mutations over the course of its evolution.

## Methods

### Sequence Collection

All sequences were collected via systematic probability sampling from NCBI Virus keyword search, optimized for ten sequences across all available entries in the query. Only full genomes were collected to avoid partially sequenced submissions. With the goal of characterizing mutations between coronavirus lineages, nine different coronaviruses were selectively chosen to represent gene mutations over time. These lineages included Human Coronavirus (str. HKU-1-12), Severe Acute Respiratory Syndrome-Related Coronavirus (hCoV-SARS str. Tor2), Middle East Respiratory Syndrome-Related Coronavirus (MERS-CoV), and Severe Acute Respiratory Syndrome Coronavirus Two (SARS-CoV-2 str. Wuhan-Hu-1, B.1.1.7, B.1.617.2, C.37, B.1.621, and C.36.3). Initially, an investigation was conducted using an n =1 for each of the species and variant groups, but proved to not provide the necessary statistical significance for a basis for analysis. An n = 10 demonstrated a statistical significance amenable to evaluation. In total, 54 whole genomes were collected, and the ORF1a and ORF1ab regions of the genome were isolated. Pp1ab coding regions for each sequence were spliced into sixteen separate respective sequences and converted to FASTA. Collectively, this data represents a total of 864 nonstructural proteins. Table 2. (right) displays each of the pp1ab domain polyproteins used for each variant.

**Table 2.**
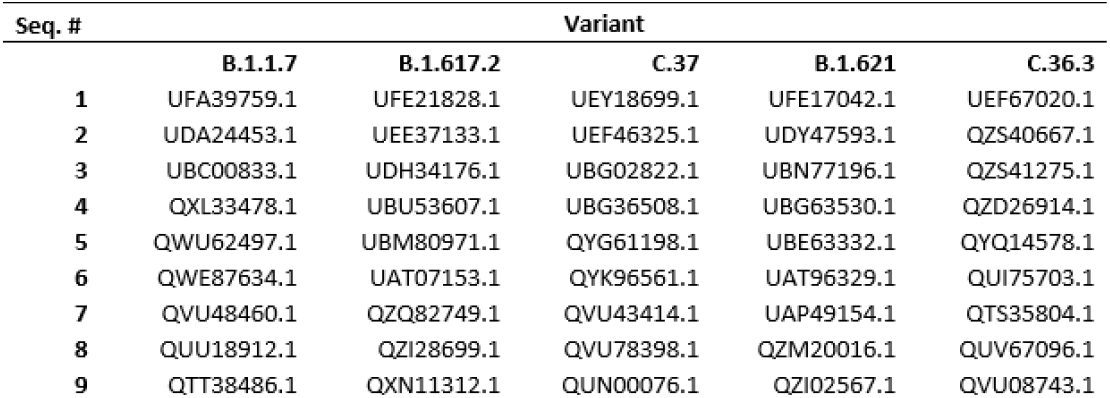
Variants and respective protein ID’s

### Alignments

A total of 18 MUSCLE alignments, 16 for each nonstructural protein, one for coronavirus family strains, and one for all 50 pp1ab sequences, were made. Alignments were done in Molecular Evolutionary Genetics Analysis (MEGA) (v. 11.0.8) using default settings. All alignments were run through Castresana Lab Gblocks Server using stringent parameters for tree construction, and headers were overwritten with Python 3.9 to a standard, *ID Strain Sample #*, format.

### Weblogo and PFAM Validation

Alignments were uploaded to UC Berkley Weblogo 2.8.2 and logos were generated for alignments with multiline symbols enabled. Logos for core replicase proteins were validated by comparison of protein family homologies on Pfam v.35. Some of these comparisons are shown in Figure 3.

### Construction of Phylogeny Trees

The alignments for all Gblocked pp1ab sequences and coronavirus families were used to generate maximum likelihood trees using MEGA (v. 11.0.8). The trees were made using the Poisson model running one hundred bootstrap replications. All other settings were default. Trees were rooted to earliest known common ancestor for the basis of analysis, and dynamic elements were added to more clearly depict the phylogenetic relationships calculated.

### Mutation Quantification and Mutagenesis Calculations

Python scripts were generated to convert .fas files to .txt formats, read into Python (v. 3.9), and compare a subject sequence to any x number of query sequences screening for independent mutation events. Independent mutation events were defined as any mutation in the amino acid sequence of collective queries occurring at any given position subtracting duplicates. Data generated from the scripts were imported to Excel in a tabular format and heatmaps for mutation count by NSP and variants were made. Calculations for average mutation count per protein and total mutation per variant across all 50 query variant sequences were made. Mutations in core replicase proteins (NSP 7,8,12) were summed and represented as a ratio of total mutagenesis to total mutations in their respective variant.

### Protein Modeling

NSP 7 and 8 predicted protein models for Wuhan-Hu-1 were made using AlphaFold 2.1.1. Structures for both of the proteins utilized standard CASP14 model parameters. MSAs generated query UniRef90 (2020), MGnify (2018), Uniclust 30 (2018) and BFD. NSP 12 predicted protein models were generated using Baker Lab deep learning model RoseTTAFold. All models were visualized in PyMol and visually compared with known crystal structures if available.

## Discussion

Phylogenetic trees generated are consistent with literature and existing knowledge on coronavirus families. It is observed from Figure 2. that SARS-CoV-2 is most closely related to SARS-CoV, with MERS-CoV evolving from a common ancestor of human coronavirus HKU-1. Weblogos shown in panels C, D, and E of Figure 3. depict homologies between the generated weblogos from the sequence alignments and known coronavirus protein families characterized in UniProt. It is reasonable to make the assumption, based on validating the web logos and the trees, that the sequences collected are accurate, and that these proteins are functional. Figure 1. Demonstrates relationships based off pp1ab protein domains across the 50 organisms sampled. It can be seen that the Alpha variant is the most conserved, followed by a mix of Lambda and Delta, then by Mu and C.36.3. These results are nearly entirely consistent with known phylogenetic relationships based on whole genome Pango lineage data. From this tree we assume that as variants become more divergent, we should observe that mutations in the core RdRp complex will represent a higher count comparative to the mutagenesis of other nonstructural proteins if the replicase is amenable to mutation as a mechanism to adapt to selective pressures in its environment.

**Figure 1.**
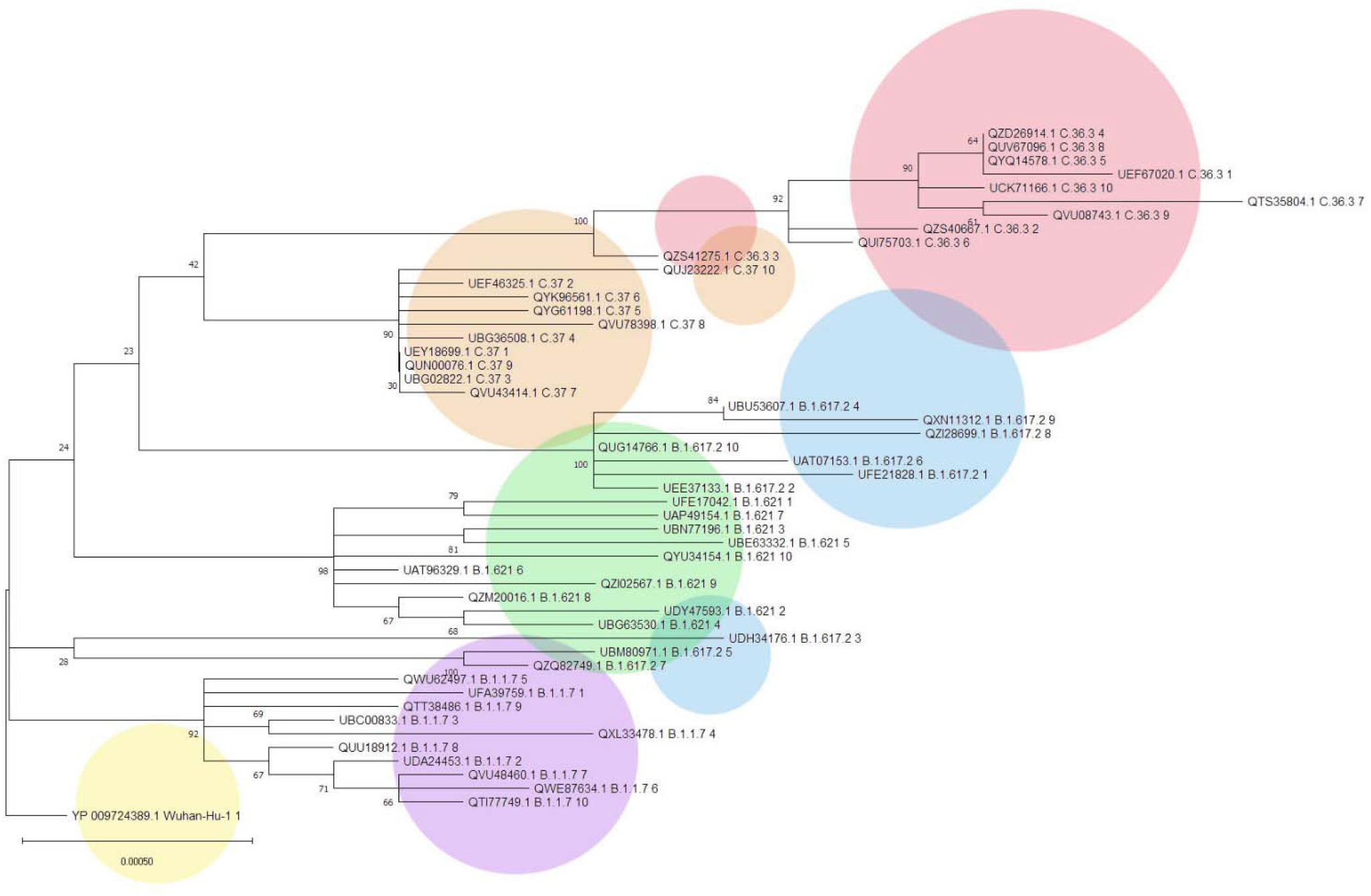
Phylogenetic tree of SARS-CoV-2 PP1ab domains from variants sampled from NCBI Virus. Re ference (Wuhan-Hu-1) shown in yellow. Alpha (B.1.1.7) (purple), Delta (B.1.617.2) (blue), Mu (B .1.621) (green), Lambda (C.37) (orange), C.36.3 (red). MEGA 11.0.8 Maximum Likelihood tree of sequences via Poisson model with 100 replication bootstrapping, all other settings default

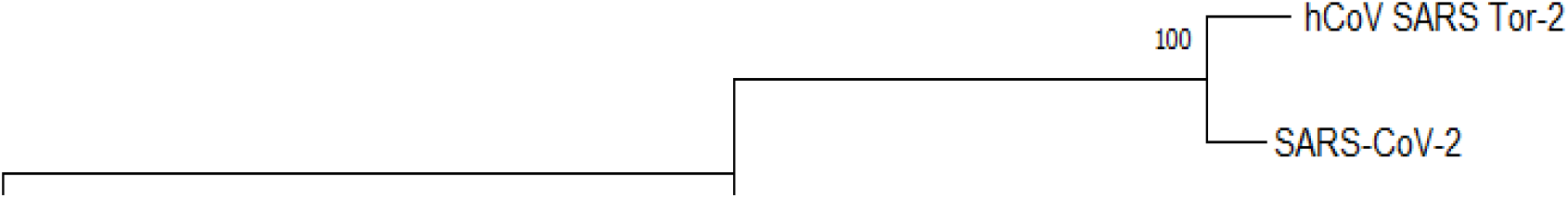

**Figure 3.**
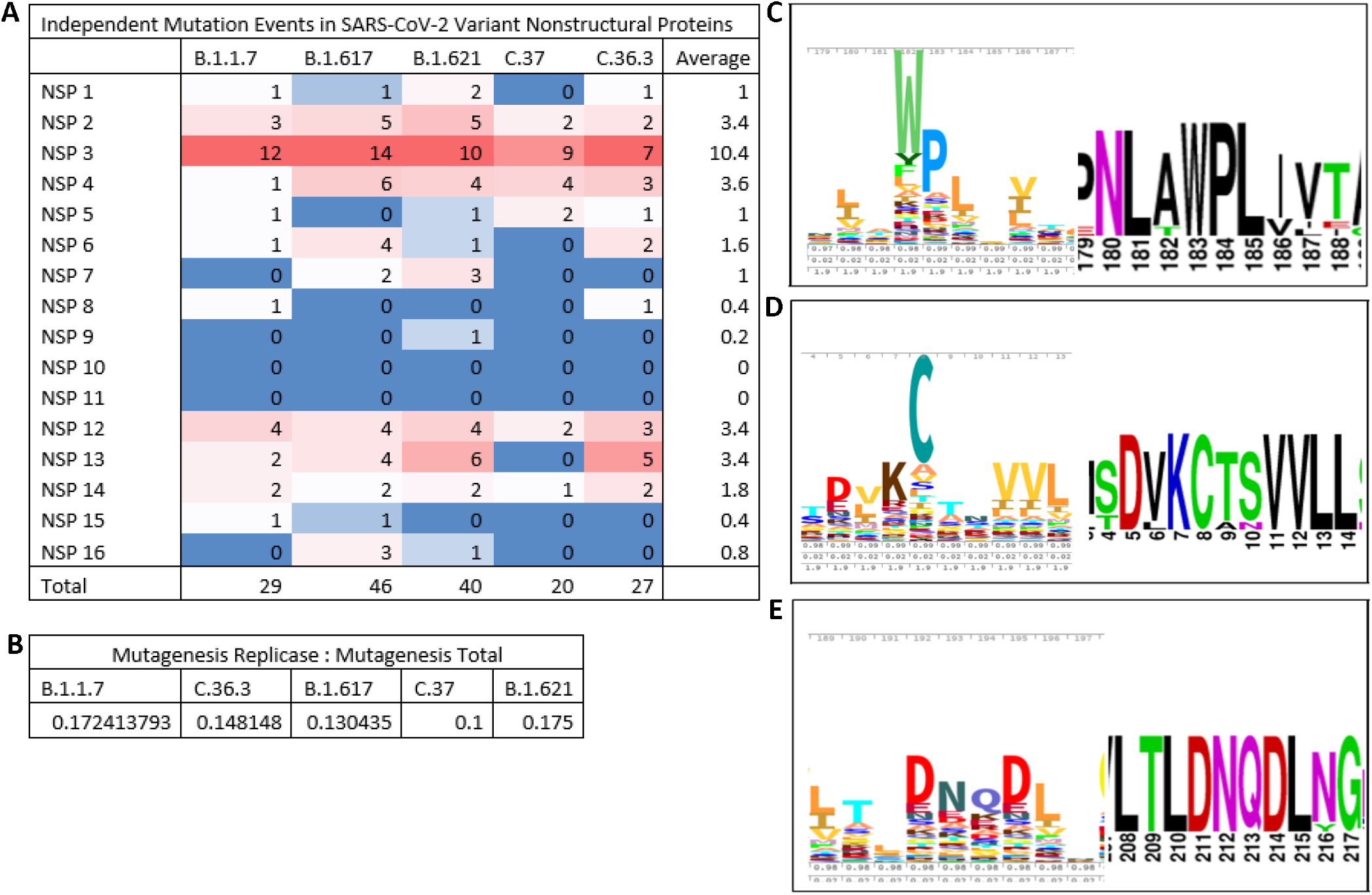
Heatmap of independent mutation events in nonstructural loci of SARS-CoV-2 variants and core NSP weblogos. Heatmap of all independent mutation events in all proteins sampled (P anel A). Mutagenesis of core replicase written as a ratio of total mutagenesis of strain (Panel B). Weblogo of NSP 7 (Panel C), NSP 8 (Panel D), and NSP 12 (Panel E). (C, D, E: Pfam left, generated l ogo right).

Figure 3’s heatmap reveals that core replicase proteins (7, 8, and 12) are among proteins with mutation counts mild to moderate compared to others. Furthermore, we can see that relative mutagenesis as a ratio of RdRp mutation to total mutation count has no observable trends that indicate that the complex has evolved in any significant way based on the sample size used. Figure 6. Shows an expected line in orange, which is a normalized average, on a scale of 0 to 0.25, of divergence based on branch length from the subject sequence expressed in Figure 1. The actual data clearly does not follow the expected trendline, and the slope of the linear regression line is 0.04, which indicates that on average there is negligible amounts of change in representation between variants.

**Figure 4.**
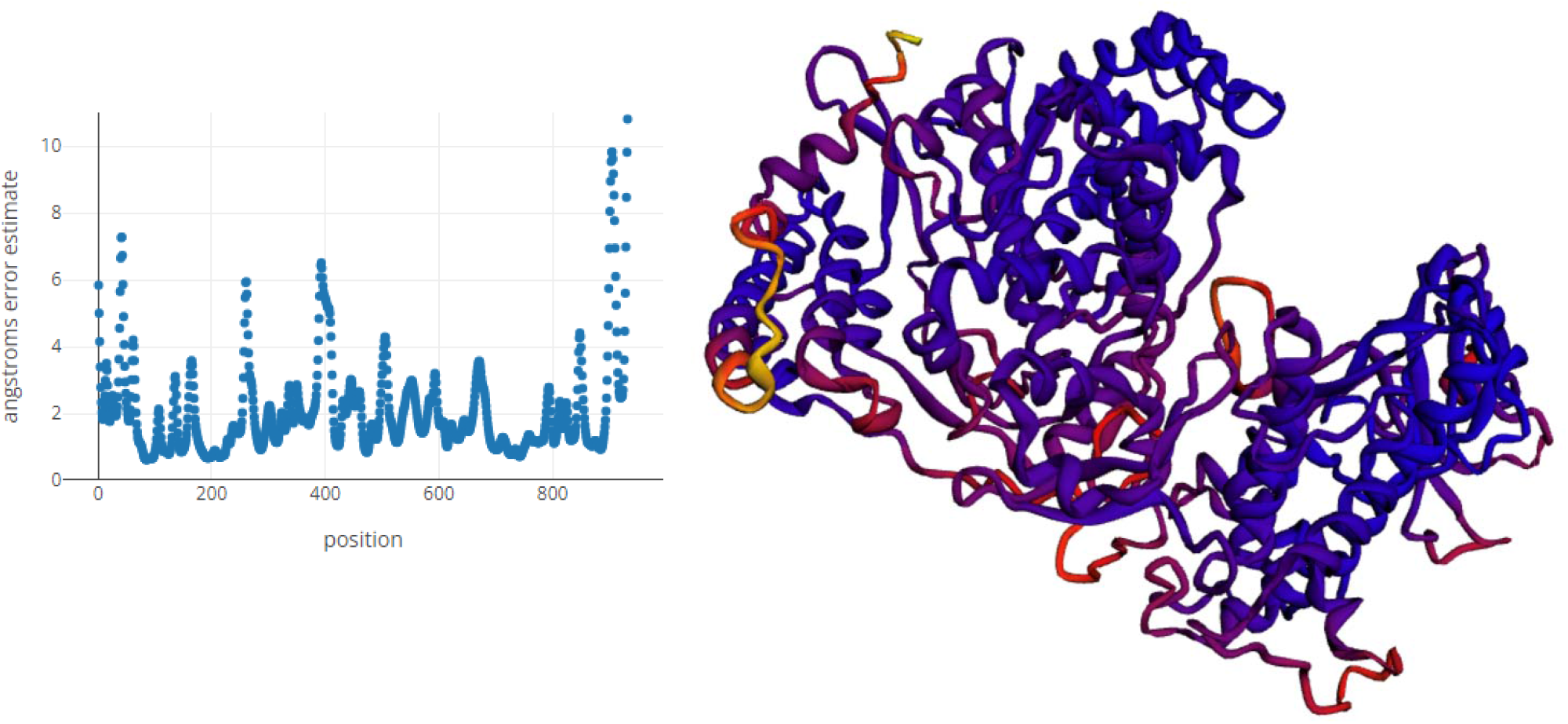
3D model of NSP 12 (RoseTTAFold 2021) Error estimate by residue (left), static capture of NSP 12 pdb (right).

**Figure 5.**
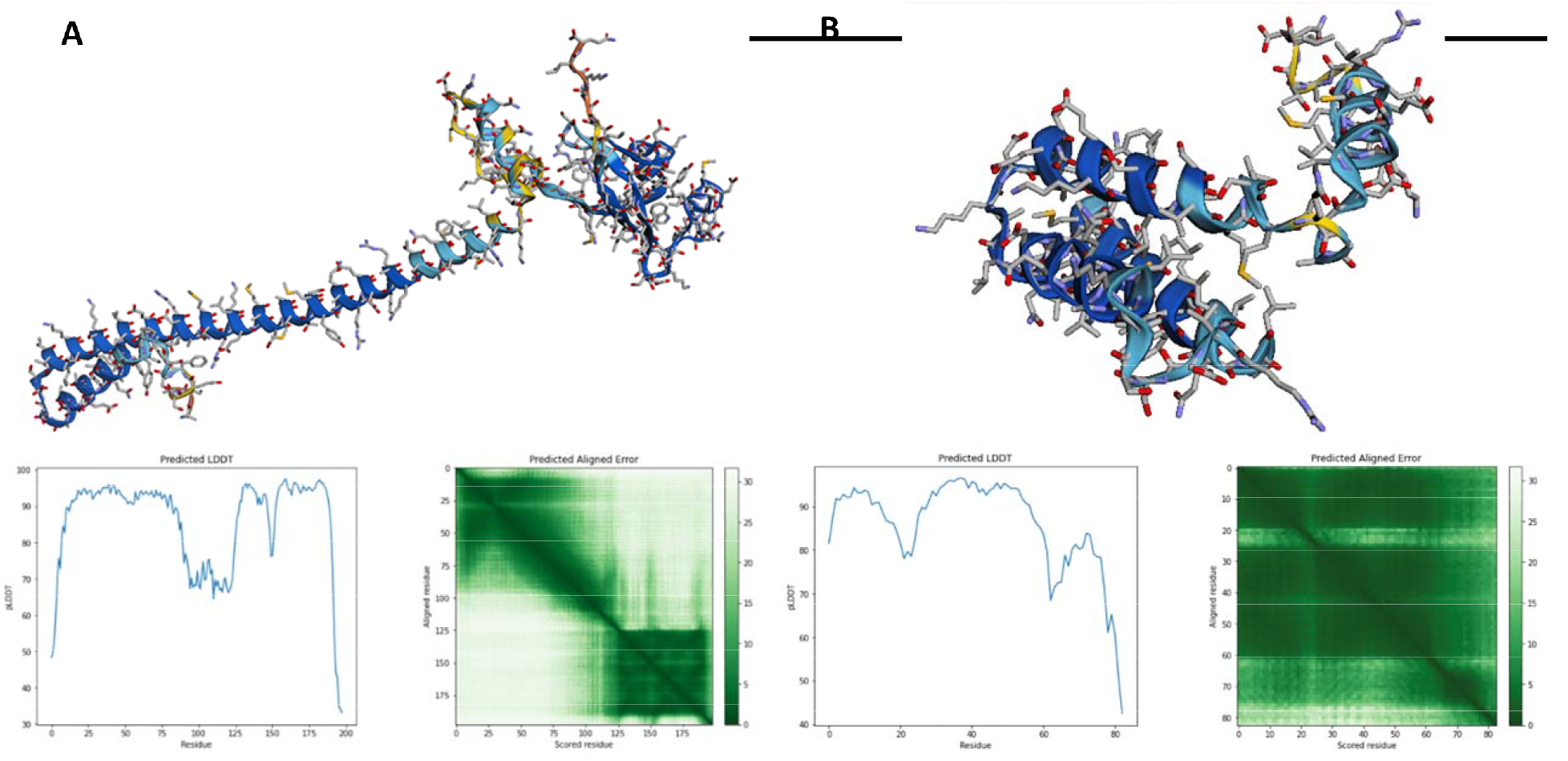
3D model of NSP 7 (Panel A) and NSP 8 (Panel B) Error estimate and alignment error estimate (below), static capture of protein models (top).

**Figure 6.**
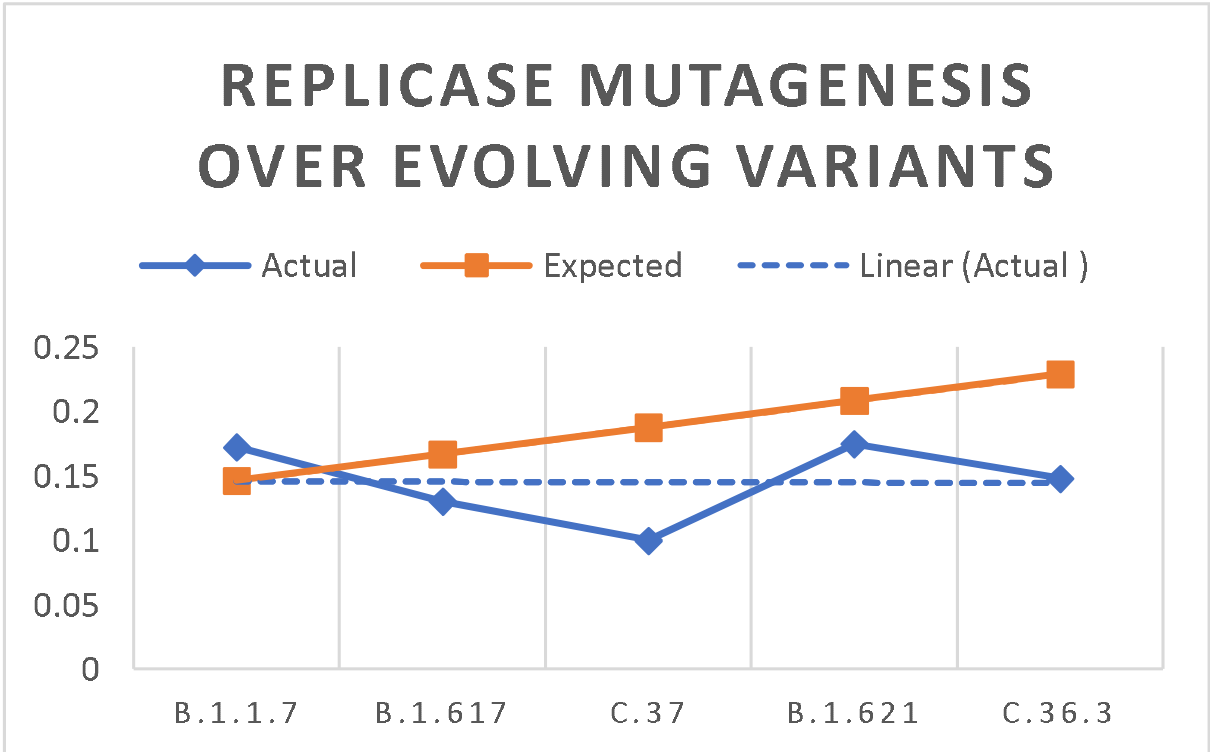
(left) Relative mutation of RdRp complex as a function of total nonstructural mutation across variants. Actual line demonstrating relative change in count of mutations. Expected line is a normalized linear estimation based off relative phylogeny as predicted by the tree.

There is a clear answer from the data collected, but there remains to be many limitations to the project which will now be addressed. Firstly, this analysis was conducted to evaluate mutation of the RdRp complex as a factor of mutation count, and not structural residue interactions as a result of mutation, and does not take into account the effect of specific chemistries in variant replicases that may potentially change the functionality of the protein. Another limitation is that only nonstructural proteins were characterized, and all accessory and structural proteins remained outside the scope of the study. It is also important to consider that reliability of conclusions may increase with the sample size. To build off the data collected, future directions should seek to ask questions to narrow scope to a single viral strain, the likely choice being a B lineage, and characterizing mutations over time in a similar manner to see exactly what changes contribute to the pathogens proliferation. It will also be helpful to see how specific chemistries change depending on these mutations, and can utilize the protein models generated from this study with known treatments such as Remdesivir.

## Conclusion

This study demonstrates with a sample size of 864 proteins across five SARS-CoV-2 variants that mutation count is most likely not a factor in viral mutagenesis of SARS-CoV-2, and suggests that the RdRp complex remains to serve as a good therapeutic target because of how highly conserved it is. This information disproves theories on mutation count going into the study, but does not provide all necessary data to conclude that the complex has not been a key factor in evolving pathogenicity. Future works should focus on the chemical interactions of mutations in the complex to characterize how they might impact the mechanism of action for therapeutics that have the potential to apply selective pressure to the virus.

## Author Approvals

All authors have seen and approved this manuscript for uploading to BioRxiv. This work has not been submitted accepted or published elsewhere. All authors declare no competing interests

## Notes

### Competing Interest Statement

The authors have declared no competing interest.

## References

1. Hillen, Hauke S., et al. “Structure of replicating SARS-CoV-2 polymerase.” Nature 584.7819 (2020): 154–156.

2. Kokic, Goran, et al. “Mechanism of SARS-CoV-2 polymerase stalling by remdesivir.” Nature communications 12.1 (2021): 1–7.

3. Helmy, Yosra A., et al. “The COVID-19 pandemic: a comprehensive review of taxonomy, genetics, epidemiology, diagnosis, treatment, and control.” Journal of clinical medicine 9.4 (2020): 1225.

4. Wu, Fan, et al. “A new coronavirus associated with human respiratory disease in China.” Nature 579.7798 (2020): 265–269.

5. Redondo, Natalia, et al. “Sars-cov-2 accessory proteins in viral pathogenesis: Knowns and unknowns.” Frontiers in Immunology 12 (2021).

6. Yadav, Rohitash, et al. “Role of Structural and Non-Structural Proteins and Therapeutic Targets of SARS-CoV-2 for COVID-19.” Cells 10.4 (2021): 821.

7. Schubert, Katharina, et al. “SARS-CoV-2 Nsp1 binds the ribosomal mRNA channel to inhibit translation.” Nature structural & molecular biology 27.10 (2020): 959–966.

8. Angeletti, Silvia, et al. “COVID□2019: the role of the nsp2 and nsp3 in its pathogenesis.” Journal of medical virology 92.6 (2020): 584–588.

9. Cornillez-Ty, Cromwell T., et al. “Severe acute respiratory syndrome coronavirus nonstructural protein 2 interacts with a host protein complex involved in mitochondrial biogenesis and intracellular signaling.” Journal of virology 83.19 (2009): 10314–10318.

10. Oostra, M., et al. “Localization and membrane topology of coronavirus nonstructural protein 4: involvement of the early secretory pathway in replication.” Journal of virology 81.22 (2007): 12323–12336.

11. Yoshimoto, Francis K. “A Biochemical Perspective of the Nonstructural Proteins (NSPs) and the Spike Protein of SARS CoV-2.” The protein journal (2021): 1–36.

12. Cottam, Eleanor M., Matthew C. Whelband, and Thomas Wileman. “Coronavirus NSP6 restricts autophagosome expansion.” Autophagy 10.8 (2014): 1426–1441.

13. Kirchdoerfer, Robert N., and Andrew B. Ward. “Structure of the SARS-CoV nsp12 polymerase bound to nsp7 and nsp8 co-factors.” Nature communications 10.1 (2019): 1–9.

14. Khan, Muhammad Tahir, et al. “Structures of SARS-CoV-2 RNA-Binding Proteins and Therapeutic Targets.” Intervirology 64.2 (2021): 55-68.1.

15. Krafcikova, Petra, et al. “Structural analysis of the SARS-CoV-2 methyltransferase complex involved in RNA cap creation bound to sinefungin.” Nature communications 11.1 (2020): 1–7.

16. Ivanov, Konstantin A., et al. “Multiple enzymatic activities associated with severe acute respiratory syndrome coronavirus helicase.” Journal of virology 78.11 (2004): 5619–5632.

17. Pillon, Monica C., et al. “Cryo-EM structures of the SARS-CoV-2 endoribonuclease Nsp15 reveal insight into nuclease specificity and dynamics.” Nature communications 12.1 (2021): 1–12.

18. Zadeh, Nadia Mohammad, et al. “Mechanism and adverse effects of COVID-19 drugs: a basic review.” International journal of physiology, pathophysiology and pharmacology 13.4 (2021): 102.

19. Gilead.” Development of Remdesivir.” Gilead, https://www.gilead.com/-/media/gilead-corporate/files/pdfs/covid-19/gilead_rdv-development-fact-sheet2020.pdf.

20. Kokic, Goran, et al. “Mechanism of SARS-CoV-2 polymerase stalling by remdesivir.” Nature communications 12.1 (2021): 1–7.

21. Li, Tingting, et al. “Phylogenetic supertree reveals detailed evolution of SARS-CoV-2.” Scientific reports 10.1 (2020): 1–9.

22. Ilmjärv, Sten, et al. “Concurrent mutations in RNA-dependent RNA polymerase and spike protein emerged as the epidemiologically most successful SARS-CoV-2 variant.” Scientific reports 11.1 (2021): 1–13.

